# The same but different: setal arrays of anoles and geckos indicate alternative approaches to achieving similar adhesive effectiveness

**DOI:** 10.1101/2020.11.03.366864

**Authors:** Austin M. Garner, Michael C. Wilson, Caitlin Wright, Anthony P. Russell, Peter H. Niewiarowski, Ali Dhinojwala

## Abstract

The functional morphology of squamate fibrillar adhesive systems has been extensively investigated and has indirectly and directly influenced the design of synthetic counterparts. Not surprisingly, the structure and geometry of exemplar fibrils (setae) have been the subject of the bulk of the attention in such research, although variation in setal morphology along the length of subdigital adhesive pads has been implicated in the effective functioning of these systems. Adhesive setal field configuration has been described for several geckos, but that of the convergent *Anolis* lizards, comprised of morphologically simpler fibrils, remains largely unexplored. Here we examine setal morphology along the proximodistal axis of the digits of *Anolis equestris* and compare our findings to those for a model gecko, *Gekko gecko*. Consistent with previous work, we found that the setae of *A. equestris* are generally thinner, shorter, and present at higher densities than those of *G. gecko* and terminate in a single spatulate tip. Contrastingly, the setae of *G. gecko* are hierarchically branched in structure and carry hundreds of spatulate tips. Although the splitting of contacts into multiple smaller tips is predicted to increase the adhesive performance of a fiber compared to an unbranched one, we posited that the adhesive performance of *G. gecko* and *A. equestris* would be relatively similar when the configuration of the setal fields of each was accounted for. We found that, as in geckos, setal morphology of *A. equestris* follows a predictable pattern along the proximodistal axis of the pad, although there are several critical differences in the configuration of the setal fields of these two groups. Most notably, the pattern of variation in setal length of *A. equestris* is effectively opposite to that exhibited by *G. gecko*. This difference in clinal variation mirrors the difference in the direction in which the setal fields of anoles and geckos are peeled from the substrate, consistent with the hypothesis that biomechanical factors are the chief determinants of these patterns of variation. Future empirical work, however, is needed to validate this. Our findings introduce *Anolis* lizards as an additional source of inspiration for bio-inspired design and set the stage for comparative studies investigating the functional morphology of these convergent adhesive apparatuses. Such investigations will lead to an enhanced understanding of the interactions between form, function, and environment of fibril-based biological adhesive systems.

## Introduction

A plethora of synthetic fibrillar adhesives inspired by the subdigital adhesive pads of some geckos have been fabricated in attempts to emulate the functional properties of their biological counterparts. Recent assessments, however, suggest that such synthetic adhesives have struggled to meet this objective (Niewiarowski et al., 2016). In light of this, Garner et al. (2019) noted that *Anolis*, a group of lizards with convergently evolved fibrillar adhesive arrays, provides an alternative model system for the study of fibrillar adhesion, observing that their structurally simpler adhesive fibrils more closely resemble the synthetic fibrils that have been manufactured. Gecko subdigital pads are composed of expanded scales (scansors) that carry arrays of structurally hierarchical fibrils (setae) terminating in hundreds of small spatulate tips (spatulae). Anoline subdigital pads are also composed of expanded scales (lamellae) that carry arrays of setae terminating in but a single large spatula. Exemplar setae of anoles have generally been described as being shorter and thinner than those of geckos (Delannoy, 2005; Maderson, 1970; Peterson, 1983; Ruibal and Ernst, 1965; Williams and Peterson, 1982). Although geckos and anoles differ in the dimensions and geometry of their adhesive setae, how these differences potentially influence adhesive performance at the level of the fiber or whole animal has received little attention.

The dimensions of a spatula, the presence of structural hierarchy, and the density of setae/spatulae are important contributors to the adhesive performance of fibrillar systems. For a single contact (i.e., spatula), larger contacts should induce greater adhesion than smaller ones (Arzt et al., 2003; Autumn et al., 2014; Johnson et al., 1971; Kendall, 1975). The concept of contact splitting predicts that the subdivision of an adhesive area (e.g., a subdigital pad) into multiple smaller contact points (e.g., setae) increases the overall adhesive force (Arzt et al., 2003). Some authors have also extended this formalism to predict that branched, hierarchical fibers should benefit further from contact splitting at multiple levels (i.e., the pad into setae and setae into multiple spatulae; Peattie and Full, 2007). Indeed, synthetic hierarchical fibers generate greater adhesion than unbranched fibers despite possessing similar contact areas (Murphy et al., 2009). Subdigital pads with greater setal/spatular density should also generate greater adhesion than those with lower setal/spatular density. Early work examining differences in the morphology of exemplar gekkotan and anoline setae found that geckos and anoles differ in many of these aspects (Peterson, 1983; Ruibal and Ernst, 1965; Stork, 1983; Williams and Peterson, 1982). Do these differences, however, translate into differences in whole animal adhesive performance between geckos and anoles? At least two studies have examined static adhesive performance in geckos and anoles and surprisingly found their adhesive systems to be functionally similar on smooth substrates (Irschick et al., 1996; Ruibal and Ernst, 1965). Thus, it is not clear what is responsible for the functional similarity in these convergent, yet morphologically distinct, adhesive systems.

In addition to the structure and geometry of individual setae, the manner in which setae are configured along the proximodistal axis of the subdigital pad impacts the effective functioning of biological fibrillar adhesives. A series of critical studies demonstrated that the morphology of gecko adhesive setae varies predictably along the length of the subdigital pad (Delannoy, 2005; Johnson and Russell, 2009; Russell et al., 2007; Russell and Johnson, 2014). Most notably, setal length increases proximodistally both within scansors and between them, and base diameter decreases proximodistally within scansors. Variation in setal length and diameter can alter the bending stiffness of the fibers, which can directly impact the contact area the fibers are capable of generating (Ge et al., 2007; Greiner et al., 2009, 2007; Johnson and Russell, 2009; Spolenak et al., 2005). The predictable variation in setal length across the subdigital pad spawned several functional hypotheses (Johnson and Russell, 2009): improvement of adhesion on rough substrates; inhibition of interference between adjacent setae during attachment; permission of simultaneous detachment of the entire battery of setae on each scansor during active distoproximal hyperextension (the subdigital pad peeling mechanism exhibited by some geckos). The subdigital pads of *Anolis*, however, detach from surfaces employing the ancestral proximodistal pattern common to lizards in general (Russell and Bels, 2001). Thus, it is possible that they exhibit different patterns of setal length if such variation is driven by differences in the biomechanics of subdigital pad peeling.

Although setal form and density have been posited to be important contributors to adhesive performance, such parameters have rarely been considered collectively when contemplating the functional similarity of anoline and gekkotan adhesive systems. This is largely because no studies have comprehensively documented the morphology and configuration of anoline setal fields, even though some work has mentioned their general variability and patterning (Peterson, 1983; Ruibal and Ernst, 1965). Exploration of these topics holds promise for enabling morphological and functional comparisons with the gecko adhesive system focused on answering outstanding questions relating to fibrillar adhesion. With this in mind, we explored setal morphology and setal field configuration of *Anolis equestris* (Merrem, 1820) and compared them, in a functional context, with similar data previously recorded for a model gecko, *Gekko gecko* (Linnaeus, 1758) (Delannoy, 2005).

## Methods

### Animals

Four *Anolis equestris* were obtained from Snakes at Sunset (Miami, FL USA). These were sacrificed via two-stage intraperitoneal injection of tricaine methanesulfonate (MS-222) (Conroy et al., 2009), fixed with 10% neutral-buffered formalin, and stored in 70% ethanol. Prior to euthanasia, anoles were cared for as described in (Niewiarowski et al., 2008). All animals were weighed before euthanasia. Snout-vent length (SVL) was measured postmortem. All protocols involving animals were approved by the University of Akron IACUC Protocol #19-07-13 NGC.

### Sample Preparation and Morphometrics

Digits of *A. equestris* were prepared similarly to the methods described by Johnson and Russell [42]. Digit IV of both the right pes and manus was removed from each specimen and sectioned parasagittally parallel to the phalanges to yield a longitudinal transect of the setal fields for imaging via scanning electron microscopy (SEM). The longitudinal sections containing phalanges were critical point dried and attached to SEM stubs with carbon tape (Fig. 1A). Samples were sputter-coated (30 seconds) with gold-palladium (Denton Vacuum Desk II; Moorestown, NJ USA) and viewed with a high-vacuum field emission SEM (JEOL 7401 FESEM; JEOL USA Inc., Peabody, MA USA).

**Figure 1.**
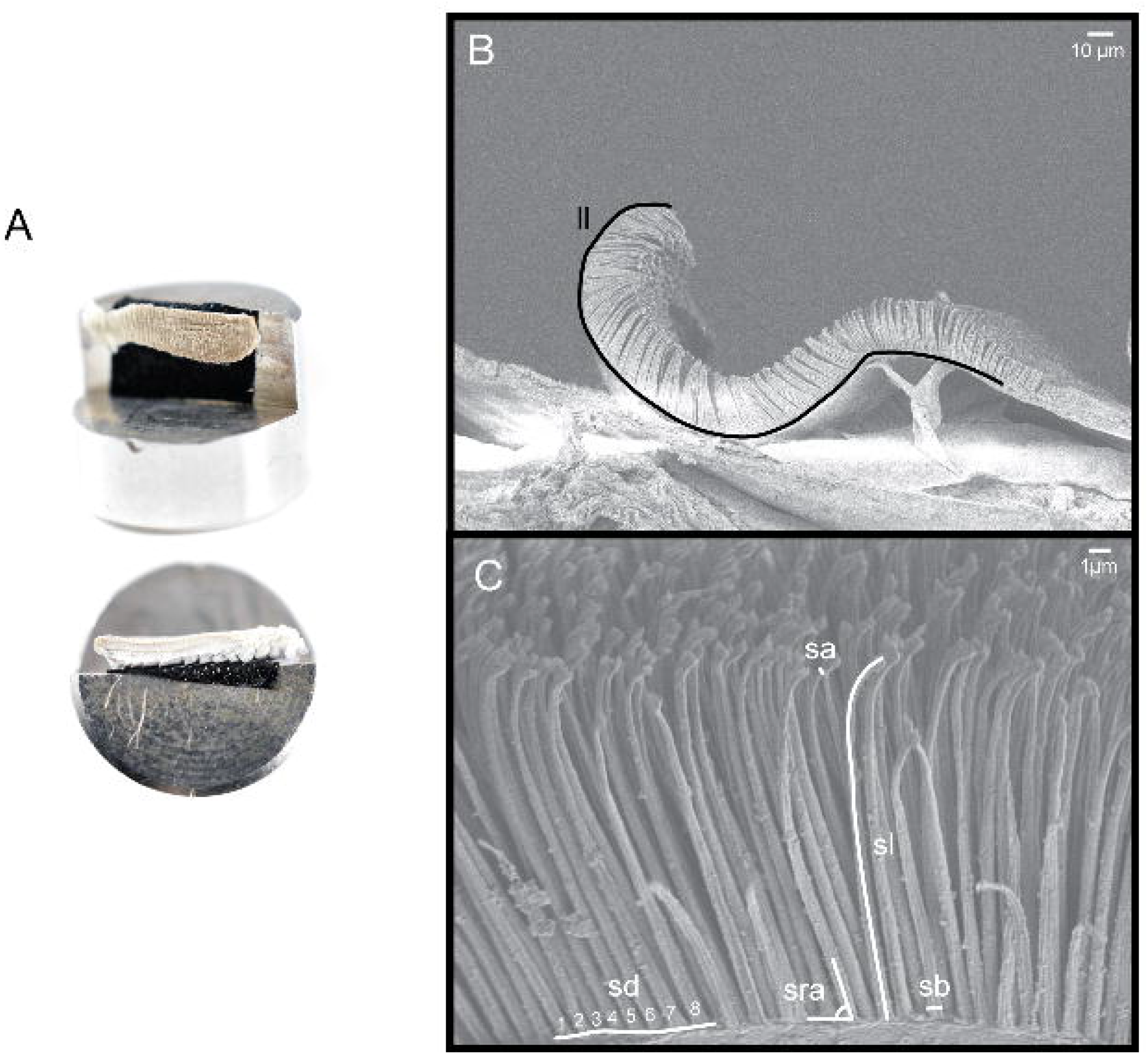
(A) Digits of *Anolis equestris* prepared for scanning electron microscopy (SEM). Digits were parasagittally sectioned, affixed to an SEM stub with carbon tape, and positioned so that the longitudinal section of the setal fields was exposed. (B) Scanning electron micrograph of an *A. equestris* lamella depicting the measurement of lamella length (ll). (C) Scanning electron micrograph of *A. equestris* adhesive setae depicting measurements of setal length (sl), setal base diameter (sb), setal apex diameter (sa), setal resting angle (sra), and setal density (sd).

ImageJ (National Institutes of Health, Bethesda, MA USA) was used to measure a variety of characteristics of lamellae and setae as seen in longitudinal section: total setal length, length of the setal stalk, setal base diameter, setal apex diameter, setal resting angle, setal density, and lamella length (Fig. 1B and 1C). Setal density was assessed by counting the number of setae along a length of a lamella and calculating an areal estimate of the number of setae; this was then extrapolated to the number of setae per mm^2^. We observed setal spacing to be similar mediolaterally, rendering our estimate of setal density accurate. Lamella length was defined as the length of the portion of the lamella bearing fully spatulate setae, which includes the epidermal free margin.

Three lamellae from each of three regions along the subdigital pad (proximal, intermediate, and distal pad regions) were imaged at ~400X to enable measurement of lamella length (Fig. 2A). In total, images of 21 distal, 15 intermediate, and 21 proximal lamellae were obtained. Images taken at ~2000X were taken along the entire proximodistal axis of these lamellae to enable measurement of setal features. Each lamella was visually subdivided into three zones (proximal, intermediate, and distal lamella zones) by splitting the total images for a lamella roughly into thirds (Fig. 2A). One image from each of these zones was selected for measurement of setal characters. Subsampling of respective areas along pad regions and lamella zones was undertaken because of the high setal densities and limitations related to sample preparation. From each image, 5-10 samples per character were examined to obtain morphometric data. Three repeat measurements were made for all setal dimensions and averaged to obtain accurate estimates. A total of 16,522 individual measurements were made.

**Figure 2.**
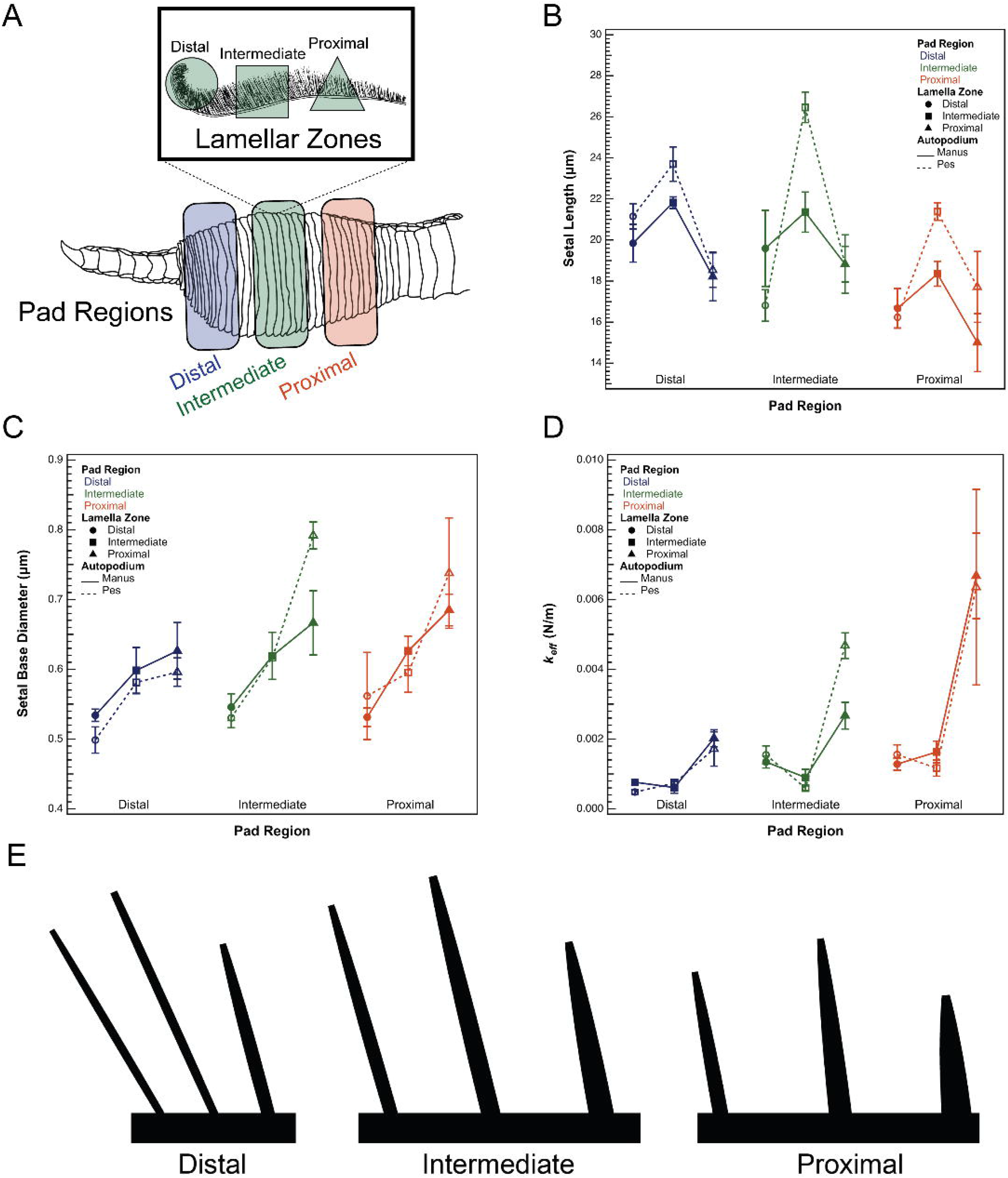
(A) Schematic depicting an *Anolis equestris* digit. Adhesive lamellae from distal (blue), intermediate (green), and proximal (orange) regions of the subdigital pad were selected for morphometric analysis. Each selected lamella was visually subdivided into three lamella zones: distal (circle), intermediate (square), and proximal (triangle). (B) Mean setal length as a function of pad region, lamella zone, and autopodium. Mean setal length increases distally along pad regions and is maximal in the intermediate lamella zone. Mean setal length is longer in the pes compared to the manus, but only in the intermediate lamella zones. Both the manus and pes exhibited similar trends in mean setal length as a function of pad region and lamella zone. (C) Mean setal base diameter as a function of pad region, lamella zone, and autopodium. Along the length of the pad of both the manus and pes, mean setal base diameter decreases proximodistally across pad regions and lamella zones. (D) Mean effective bending modulus (*k*_*eff*_) as a function of pad region, lamella zone, and autopodium. Mean *k*_*eff*_ decreases proximodistally along pad regions and lamella zones. (E) The general morphological trends of the setal field configuration of *Anolis equestris* found in this study. Figure not drawn to scale.

### Estimation of effective bending stiffness of anoline setae

Collection of anoline setal morphometric data allowed the effective bending stiffness of anoline setae to be estimated. We found these setae to be noticeably tapered, requiring the modification of the traditional equation used to estimate bending stiffness of cylinders with fixed radius because the moment of inertia varies along the length of the fiber (Caliaro et al., 2013). This modification is simply the multiplication of the bending stiffness of a fixed radius cylinder (*k*) by what we call the tapering ratio (*t*), which is the ratio of the apex radius (*R*_*a*_) to the base radius (*R*_*b*_). Thus, the effective bending stiffness (*k*_*eff*_) of an anoline seta can be calculated as

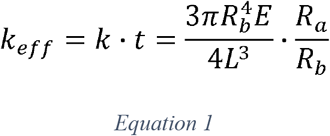

where *E* is Young’s modulus of β-keratin (~1 GPa) and *L* is the length of the setal stalk. It is evident that more drastically tapered structures (*t* ≈ 0) will have a significantly reduced effective bending stiffness. We use the above estimation to examine effective bending stiffness (*k*_*eff*_) of setae along the proximodistal axis of anoline subdigital pads.

### Statistical analyses

Setal morphometric data obtained from one high magnification image were averaged to obtain mean setal morphometrics for that lamella zone. Mean effective setal bending stiffness *k*_*eff*_ was calculated based on mean morphometric values.

Prior to robust statistical analysis, the overall mean values of each of the setal morphological characters for each individual were regressed against the body mass and SVL of that individual using ordinary least squares (OLS) regression. There were no significant linear relationships between body size and any of the morphological parameters (P > 0.05), permitting non size-corrected data to be used in the analyses.

Analyses of variance (ANOVA) were used to compare setal morphometric and *k*_*eff*_ data as a function of pad region, lamella zone, autopodium (manus or pes), and all possible interactions. Interactions that were not significant after an initial analysis were removed. In the case of significant effects (P < 0.05), post-hoc Tukey’s Honest Significant Difference (HSD) tests were performed to determine where particular significant differences occurred. In the case of setal density and *k*_*eff*_, data were log-transformed to meet the homogeneity of variance and normality assumptions of ANOVA. Two variables, setal base diameter and setal resting angle, violated one of the assumptions of ANOVA and transformations were unsuccessful in alleviating this, requiring the use of nonparametric statistical methods. Variance between lamella zones was heterogeneous for setal base diameter, thus Welch’s ANOVA was used to determine differences in base diameter as a function of pad region, lamella zone, or autopodium. In the case of significant factors, the Games-Howell nonparametric post hoc test was used to determine which groups were significantly different from one another. The residuals of setal resting angle were not normally distributed, thus the Wilcoxon rank sum test was used to examine whether setal resting angle varied significantly as a function of pad region, lamella zone, or autopodium. In the case of significant factors, nonparametric pairwise comparisons using the Wilcoxon method were performed post hoc and alpha values corrected using the sequential Bonferroni method. All statistical tables are included in the Supporting Information for reference.

## Results

### *Setal morphology of* Anolis equestris

The setal morphology of *A. equestris* varies considerably across the subdigital pad. Setae range in length from 6.56 to 29.98 μm (Fig. 2B), with base diameters ranging from 241 nm to 1.25 μm (Fig. 2C). The setae of *A. equestris* are markedly tapered, with apex diameters generally being smaller than the corresponding base diameter. Setal apex diameters range from 98 to 603 nm (SI Fig. 1). Setal resting angle varies from 28.27 to 130.15° (SI Fig. 2). Setal density ranges from 5.68 × 10^5^ to 2.20 × 10^6^ setae mm^−2^ (SI Fig. 3). The length of lamellae varies from 93.37 to 270.80 μm (SI Fig. 4).

### *Setal field configuration of* Anolis equestris

Setal length varies significantly as a function of pad region (DF = 2, F = 15.60, P < 0.0001), lamella zone (DF = 2, F = 25.77, P < 0.0001), autopodium (DF = 1, F = 6.31, P = 0.015), and the interaction between lamella zone and autopodium (DF = 2, F = 3.39, P = 0.042; Fig. 2B, SI Tables 1, 5, and 10). For both autopodia, setae from the intermediate and distal pad region are significantly longer than those from the proximal pad region (distal vs. proximal: P < 0.0001; intermediate vs. proximal: P = 0.0002), but setae of the distal and intermediate pad region are not significantly different in length (P = 0.962). Setae are longest in the intermediate lamella zone (intermediate vs. distal and intermediate vs. proximal: P < 0.0001), but setae of the distal and proximal zones are not significantly different in length (P = 0.40). Setae of the intermediate lamella zones of the pes are significantly longer than those of the manus (P = 0.014; Fig. 2B). Trends in setal length between pad regions are identical for the manus and pes. Trends in setal length between lamella zones are similar for the manus and pes, although setae from the distal lamella zone are not significantly different in length than setae from the intermediate lamella zone in the manus (P = 0.151).

Welch’s ANOVA revealed highly significant variation in setal base diameter depending on lamella zone (DF = 2, F = 23.02, P < 0.0001) and marginally non-significant variation across pad regions (DF = 2, F = 3.31, P = 0.05; Fig. 2C, SI Tables 1, 6, and 11). There were no significant differences in setal base diameter as a function of autopodium (P = 0.92). We observed significant decreases in setal base diameter proximodistally across lamella zones (all pairwise comparisons P < 0.05), and a general trend of decreasing setal base diameter proximodistally along pad regions.

Setal apex diameter varies significantly as a function of autopodium (DF = 1, F = 22.47, P < 0.0001), pad region (DF = 2, F = 8.13, P = 0.0009), lamella zone (DF = 2, F = 7.27, P = 0.0018), and the interaction between pad region and lamella zone (DF = 4, F = 3.29, P = 0.018; SI Fig. 1, SI Tables 1, 5, and 12). Setal apex diameter is significantly greater for setae of the pes than those of the manus (P < 0.0001). Setal apex diameter initially appears to decrease proximodistally between pad regions and lamella zones, but examination of the significant interaction term revealed that this was not the case. Setal apex diameter is significantly smaller in distal pad regions compared to proximal ones, but only for the proximal lamella zone (P = 0.0003). There are no other significant differences in setal apex diameter between pad regions. In the proximal pad region, setal apex diameter is significantly smaller in the distal lamella zones compared to the proximal lamella zones (P = 0.0003). Setae of the intermediate lamella zones of the proximal pad region do not, however, vary significantly in setal apex diameter from the distal (P = 0.504) or proximal lamella zones (P = 0.111). Setae do not vary significantly in apex diameter along lamella zones in the intermediate and distal pad regions.

Effective setal bending stiffness (*k*_*eff*_) varies significantly as a function of pad region (DF = 2, F = 23. 07, P < 0.0001) and lamella zone (DF = 2, F = 50.65, P < 0.0001; Fig. 3A, SI Tables 2, 7, and 13). Setal *k*_*eff*_ does not differ significantly between autopodia (DF = 1, F = 0.6, P = 0.44). Across pad regions, setal *k*_*eff*_ becomes significantly smaller proximodistally (all pairwise comparisons P < 0.05). Setal *k*_*eff*_ of distal and intermediate lamella zones is significantly less than that of the proximal lamella zones (both comparisons P < 0.0001), whereas setal *k*_*eff*_ of distal and intermediate lamella zones are not significantly different (P = 0.0632).

**Figure 3.**
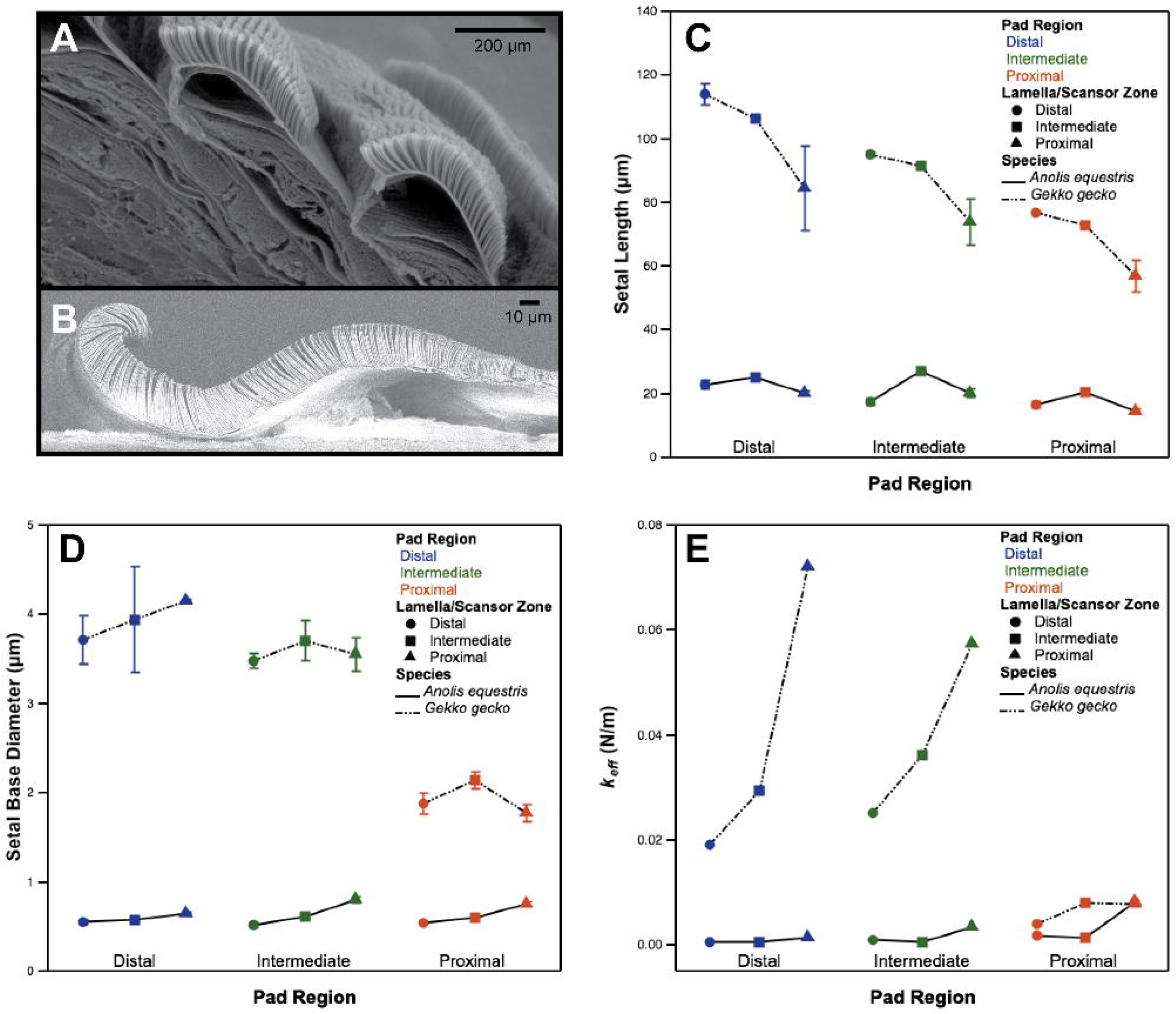
Comparative setal morphology of *Anolis equestris* and *Gekko gecko*. (A) Representative scanning electron micrograph of two scansors from the setal field of *G. gecko*, from Delannoy (2005). (B) Representative scanning electron micrograph of a lamella from the setal field of *A. equestris*. (C) Setal length of *A. equestris* and *G. gecko* setae as a function of pad region and lamella/scansor zone. (D) Setal base diameter of *A. equestris* and *G. gecko* setae as a function of pad region and lamella/scansor zone. (E) Effective bending stiffness (*k*_*eff*_) of *A. equestris* and *G. gecko* setae as a function of pad region and lamella/scansor zone.

A Wilcoxon rank sum test revealed that setal resting angle varies significantly as a function of pad region (ChiSquare = 6.321, DF = 2, P = 0.042) and lamella zone (ChiSquare = 20.065, DF = 2, P < 0.0001; SI Fig. 2, SI Tables 2, 8, and 14), but not for equivalent regions of the autopodia (P = 0.16). Resting angle is significantly lower in the distal pad region than the proximal pad region (P = 0.01), yet setae of the intermediate pad region do not differ significantly in resting angle when compared with the proximal or distal pad regions (intermediate vs. proximal: P = 0.248, intermediate vs. distal: P = 0.369). Setal resting angle decreases significantly proximodistally amongst lamella zones (all pairwise comparisons P < Bonferonni adjusted alpha ⍰).

ANOVA revealed that setal density does not vary significantly as a function of pad region, lamella zone, limb, or any of their interactions (whole model: DF = 5, F = 5.016, P = 0.11; SI Fig. 3 and SI Table 3).

Lamella length varies significantly along pad regions and between the manus and pes (P = 0.015 and 0.0009, respectively; SI Fig. 4, SI Tables 4, 9, and 15). Lamellae of the distal pad region are significantly shorter than those of the proximal and intermediate pad region (distal vs. proximal: P = 0.041; distal vs. intermediate: P = 0.022), whereas the length of lamellae of the intermediate pad region are not significantly different from that of lamellae of the proximal pad region (P = 0.985). Lamellae of the pes are significantly longer than those of the equivalent regions of the manus (P = 0.003).

## Discussion

### *Setal field configuration of* Anolis equestris – *an overview*

Many setal morphological features of *A. equestris* vary predictably along the proximodistal axis of the subdigital pad (Fig. 2E). The general trends are as follows. Overall, setal length increases proximodistally along pad regions, but is maximal in intermediate lamella zones. Setal base diameter and setal resting angle decreases proximodistally along pad regions and lamella zones. Lamella length decreases markedly proximodistally along the subdigital pad. As a consequence of variation in setal length and setal base diameter, effective setal bending stiffness (*k*_*eff*_) decreases significantly along the proximodistal axis of both the subdigital pad and lamella zones, indicating that setae of the distal regions of the subdigital pad and lamellae are considerably more flexible than those situated more proximally. Setal length of intermediate lamella zones is significantly greater for the pes than the manus but there are no significant differences between the effective bending stiffness (*k*_*eff*_) of the setae of the manus and pes.

### *Comparative morphology of* Anolis equestris *and* Gekko gecko *setae and its functional implications*

With regard to fibrillar adhesion, the Tokay gecko (*Gekko gecko*) is arguably the most frequently studied gekkonid. This makes it an excellent candidate for morphological comparison with a similarly-sized species of *Anolis*. Comparing data for a representative individual of *Gekko* and *Anolis* from our present study, the setae of *A. equestris* are 3-5 times shorter and 2-7 times narrower at their base than those of *G. gecko* (Fig. 4B and 4C). Setal resting angle in *A. equestris* ranges from being similar to that of *G. gecko* to nearly twice as great, and setal density of *A. equestris* is between 40-88 times greater than that for *G. gecko*. Although we had difficulty reliably obtaining images of the spatulae of *A. equestris* along the proximodistal axis of the subdigital pad, they appear to be approximately 0.75 μm wide, and are thus likely between 3-9 times greater in width than those of *G. gecko*.

**Figure 4.**
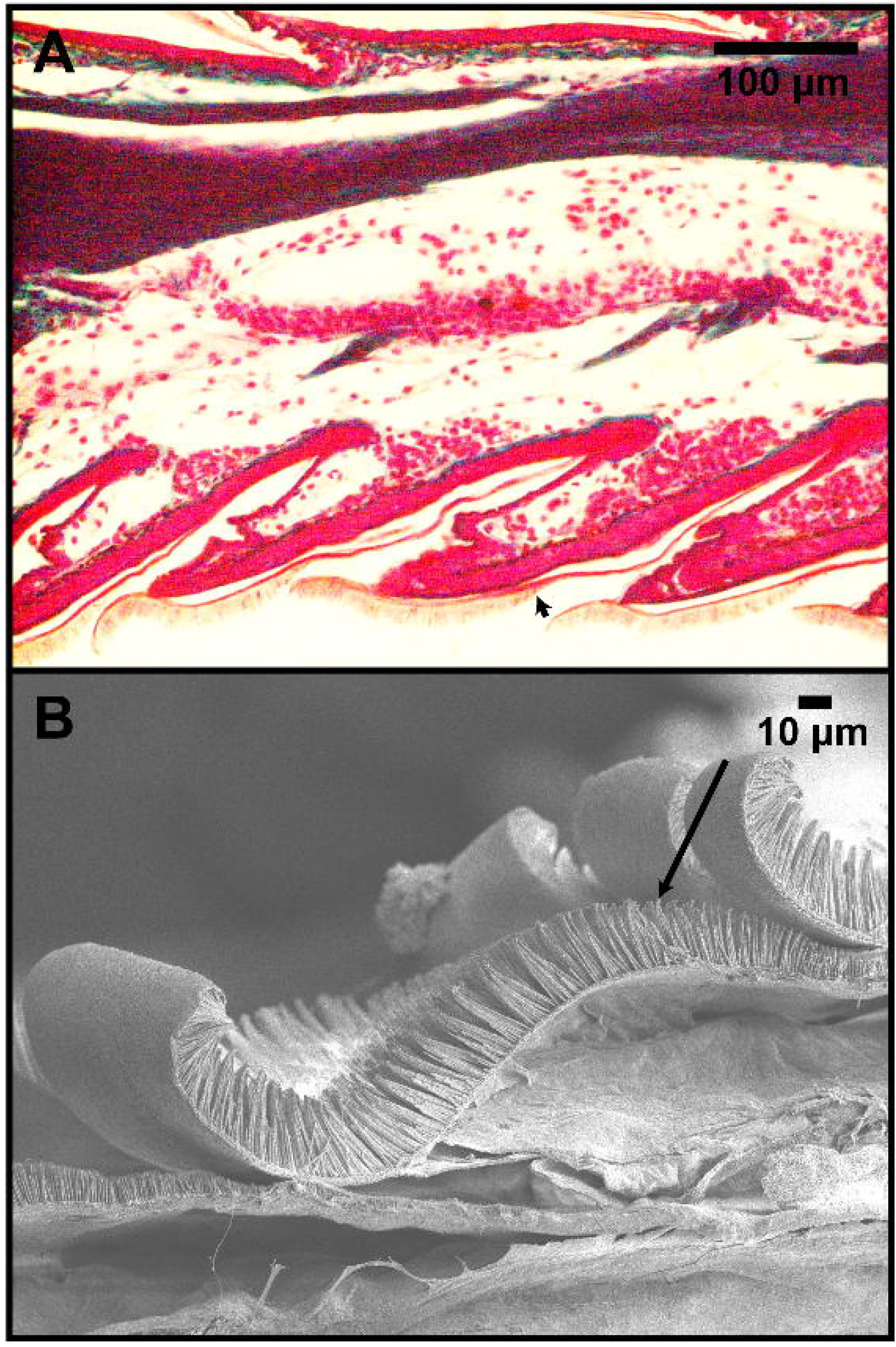
The overlap of anoline setae of the proximal lamella zone by those of the distal lamella zone of the next most proximal lamella in (A) a histological section of the subdigital pad of *Anolis sagrei* (Cocteau in A.M.C Duméril and Bibron, 1837) and (B) a scanning electron micrograph of *A. equestris* lamellae. Arrows indicate the approximate location where the overlap of the two lamellae ends when pressed in contact with the substrate.

Although both geckos and anoles have greater tip densities than other organisms employing fibrillar adhesion (Arzt et al., 2003; Garner et al., 2019; Ruibal and Ernst, 1965; Williams and Peterson, 1982), the setae of *G. gecko* and *A. equestris* differ in terms of their overall morphology. A single *G. gecko* seta terminates in hundreds of spatulae (Delannoy, 2005), whereas that of *A. equestris* carries only one. Thus, the setae of *G. gecko* exhibit contact splitting, consistent with the idea that the subdivision of a single fiber should increase its adhesive force potential. If the only difference between the setae of geckos and anoles was the presence of structural hierarchy, the magnitude of gecko adhesion would certainly be greater than that of anoles, as has been demonstrated for synthetic fibrillar systems (Murphy et al., 2009). The data collected here, however, demonstrate that gecko and anole setae not only vary in their hierarchical structure, but also in their density, setal size, and spatula width. Several authors (Arzt et al., 2003; Autumn et al., 2014, 2002; Huber et al., 2005; Spolenak et al., 2005) have modeled spatulae as hemispheres to employ the Johnson-Kendall-Roberts contact theory of elastic solids to estimate the pull-off force of a single spatula (see SI). Using this formalism, we estimated that a single spatula of *Anolis equestris* with a radius of 0.375 μm is capable of generating 0.09 μN of pull-off force, assuming a work of adhesion (*γ*) typical of van der Waals interactions (50 mJ m^−2^). When multiplied by the range of setal densities observed in this study, this results in 0.09-0.14 N mm^−2^ of normalized adhesive force. Using the range of spatula diameters and densities measured by Delannoy (Delannoy, 2005), *G. gecko* is estimated to generate between 0.05-0.25 N mm^−2^ of normalized adhesive force. The predicted adhesive force of *A. equestris* falls well-within the range of that estimated for *G. gecko*. Thus, our theoretical calculations indicate that the benefit of structural hierarchy exhibited by individual *G. gecko* setae does not translate into overall greater adhesive performance compared to *A. equestris* because of the greater density of spatulae present on the subdigital pads of *A. equestris* compared to *G. gecko*. (Irschick et al., 1996). These findings provide a potential morphological explanation for the similarity in whole organism adhesive performance between anoles and geckos on smooth substrates (Irschick et al., 1996; Ruibal and Ernst, 1965). Of course, structural hierarchy has been implied to influence a number of other properties of fibrillar systems, including self-matting, fiber fracture, flaw tolerance, and adhesion to rough substrates (Persson, 2003; Spolenak et al., 2005; Yao and Gao, 2006). Thus, any functional consequences of the absence of structural hierarchy of anoline setae may become more apparent in dynamic contexts or under less-than-ideal conditions (e.g., rough and undulant substrates).

### Setal field configuration in anoline and gekkonid adhesive apparatuses – functional implications seen through a comparative lens

Historically, most authors have focused on the geometry and characteristics of a single exemplar seta and how this, by extrapolation, relates to the function of the entirety of the adhesive apparatuses of squamates. However, setal variation along the subdigital pad and individual lamellae also has the potential to influence the performance and functional properties of fibrillar adhesive systems (Johnson and Russell, 2009; Russell and Johnson, 2014, 2007). In light of this, we turn from consideration of comparisons of setal morphology of *A. equestris* and *G. gecko* to comparisons of the configuration of their setal fields.

Although having some similarities, the setal field configuration of *A. equestris* and *G. gecko* is quite different. Most strikingly, the setae of *A. equestris* are longest in the intermediate lamella zone, whereas in *G. gecko*, setae increase progressively in length proximodistally along scansor zones. Examination of histological sections from other species of *Anolis* and SEM micrographs of the lamellae of *A. equestris* (Fig. 4) demonstrate that the shorter proximal setae may be covered by the dorsal aspect of the epidermal free margin of the immediately preceding (i.e., more proximal) lamella when the subdigital pad is placed in contact with the substrate. Thus, the setal field exposed to and in contact with the substrate exhibits decrease in setal length from intermediate portions of lamellae to those situated more distally, effectively opposite to the trend observed in *G. gecko*. Trends in setal base diameter are also markedly different. In *A. equestris*, setal base diameter decreases proximodistally along the zones of the lamellae and in succeeding pad regions, whereas in *G. gecko*, setal base diameter increases proximodistally along pad regions. From one scansor zone of *G. gecko* to the next, there is little proximodistal variation in setal base diameter, although it appears to be maximal in the intermediate scansor zone of the proximal pad region.

Variability in the trends of setal length and diameter between geckos and anoles has the potential to influence variation in their material properties along the subdigital pad. For example, the proximodistal increase in gecko setal length is thought to reduce bending stiffness along scansors. As such, setae situated more distally on a scansor may be able to achieve greater substrate contact and induce greater adhesive force than those that are more proximally situated (Johnson and Russell, 2009). Although there are marked differences between trends in setal length and diameter along scansors/lamellae in *G. gecko* and *A. equestris*, trends in the effective bending stiffness (*k*_*eff*_) of the setae of these two species are similar (i.e., there is a general decrease in *k*_*eff*_ proximodistally along each lamella and along the digit as a whole). Furthermore, the high aspect ratio of biological fibrillar adhesives is thought to considerably diminish the effective elastic modulus (*E*_*eff*_) of fibril arrays to that within the range of tacky materials (Autumn et al., 2006). Employing the formalism introduced by Autumn et al. (Autumn et al., 2006) with slight modifications to account for tapering (see SI for details), we found that the average *E*_*eff*_ of anoline setal arrays is estimated to range between 50.4 kPa – 2.9 MPa along the proximodistal axis of the subdigital pad. Although the dimensions and configuration of anoline setae (length, diameter, resting angle, and density) result in a wide range of *E*_*eff*_, eight out of nine regions examined have *E*_*eff*_ less than 2 MPa, which is within the range of tacky materials used for pressure-sensitive adhesives (e.g., Sylgard 184, polydimethylsiloxane (Bartlett et al., 2012; Khanafer et al., 2008)). As such, the major differences between gecko and anole setal field configuration likely have little, if any, consequences for differential contact and adhesion along the proximodistal axis of the subdigital pad. There are, however, a number of higher-order anatomical differences between the subdigital adhesive systems of geckos and anoles (Russell, 2017; Russell and Gamble, 2019). How these may interact with setal morphology and setal field configuration in relation to their effect on adhesion is unknown.

Setal field configuration also has the potential to impact other properties of fibrillar adhesive systems beyond their material properties. Setal length variation along gecko scansors, for example, has been implicated in permitting effective release of subdigital pads during distoproximal hyperextension (distal-to-proximal subdigital pad peeling exhibited by some geckos) (Johnson and Russell, 2009). Many geckos employ distoproximal peeling of their subdigital pads from the substrate and it has been calculated that the graded variation in setal length along the proximodistal axis of each scansor is consistent with all setae of that single scansor attaining their critical angle of detachment simultaneously, as opposed to successively for each setal row (Johnson and Russell, 2009). *Anolis* lizards, however, peel their digits proximodistally (Russell and Bels, 2001). The patterning of setal length in *Anolis equestris* is virtually opposite that observed in geckos and is thus consistent with the simultaneous release hypothesis, although more empirical work is needed to corroborate this. Among geckos, some genera may employ proximodistal hyperextension, but comprehensive investigations of their locomotor kinematics and setal field configuration are lacking (Russell and Gamble, 2019). Furthermore, variation in bending stiffness could also influence setal release; setae that are less stiff should reach their critical detachment angles more easily than stiffer ones. Thus, studies of setal field configuration and subdigital pad biomechanics in broader phylogenetic contexts, in addition to those utilizing model synthetic adhesives, are critical for the further exploration of these topics.

## Conclusions

*Anolis* lizards have long been considered a model system for the study of adaptive radiation and evolutionary key innovations (Losos, 2011, 1994), yet the microstructure of their adhesive subdigital pads, one of two proposed key innovations (Losos, 2011), have remained relatively understudied. Herein we report on our comprehensive examination of the setal morphology and setal field configuration of a model crown giant anole, *Anolis equestris*, and make comparisons with similar data collected for a model gecko, *Gekko gecko* (Delannoy, 2005). The setae of *A. equestris* are generally thinner, shorter, and present in higher densities than those of *G. gecko*. Although contact splitting is predicted to increase the adhesive performance of a branched fiber compared to an unbranched one, we discovered that, by taking into account the configuration of their entire setal fields, the predicted adhesive performance of *G. gecko* and *A. equestris* setal fields is relatively similar. Anoles compensate for the lack of branched setae by carrying more setae per unit area. These findings highlight the importance of incorporating morphological variability into functional hypotheses, extrapolations, and analyses. Furthermore, we found that the pattern of variation of setal length is effectively opposite in *A. equestris* to that of *G. gecko*. Considering this difference in direction of clinal variation mirrors the difference in the direction in which the setal fields of geckos and anoles are peeled from the substrate, our findings are consistent with the hypothesis that biomechanical factors are the chief determinants of these patterns of variation. Future empirical work, however, is needed to validate this hypothesis. Our work here sets the stage for further comparative studies that explore the functional differences between two convergent squamate adhesive systems and the connections between form, function, and environment as they relate to biological fibrillar adhesive systems.

## Acknowledgments

We thank Alexandra Tomasko for assistance in collecting preliminary data for this project and Dr. Henry Astley for additional specimens. We also thank Dr. Bojie Wang for assistance with scanning electron microscopy. Finally, we acknowledge the helpful support and insight of the Gecko Adhesion Research Group and the Dhinojwala Research Group.

## Author contributions

A.M.G. and M.C.W. designed the initial study; A.M.G., M.C.W., A.P.R., P.H.N., and A.D., refined and finalized the study design; A.M.G., M.C.W., and C.W. collected the study data; A.M.G. statistically analyzed the data and generated figures/tables; A.M.G., M.C.W., and C.W. drafted initial versions of the manuscript; All authors revised the manuscript and approved the final version for publication.

## Data availability statement

Data used in this study are available in the figshare repository at: https://doi.org/10.6084/m9.figshare.12637019.

## References

Arzt, E., Gorb, S., Spolenak, R., 2003. From micro to nano contacts in biological attachment devices. Proceedings of the National Academy of Sciences of the United States of America 100, 10603–10606. https://doi.org/10.1073/pnas.1534701100

Autumn, K., Majidi, C., Groff, R.E., Dittmore, A., Fearing, R., 2006. Effective elastic modulus of isolated gecko setal arrays. Journal of Experimental Biology 209, 3558–3568.

Autumn, K., Niewiarowski, P.H., Puthoff, J.B., 2014. Gecko adhesion as a model system for integrative biology, interdisciplinary science, and bioinspired engineering. Annual Review of Ecology, Evolution and Systematics 45, 445–470. https://doi.org/10.1146/annurev-ecolsys-120213-091839

Autumn, K., Sitti, M., Liang, Y.A., Peattie, A.M., Hansen, W.R., Sponberg, S., Kenny, T.W., Fearing, R., Israelachvili, J.N., Full, R.J., 2002. Evidence for van der Waals adhesion in gecko setae. Proceedings of the National Academy of Sciences of the United States of America 99, 12252–12256.

Bartlett, M.D., Croll, A.B., King, D.R., Paret, B.M., Irschick, D.J., Crosby, A.J., 2012. Looking beyond fibrillar features to scale gecko◻like adhesion. Advanced Materials 24, 1078–1083.

Caliaro, M., Schmich, F., Speck, T., Speck, O., 2013. Effect of drought stress on bending stiffness in petioles of *Caladium bicolor* (Araceae). American Journal of Botany 100, 2141–2148. https://doi.org/10.3732/ajb.1300158

Conroy, C.J., Papenfuss, T., Parker, J., Hahn, N.E., 2009. Use of tricaine methanesulfonate (MS222) for euthanasia of reptiles. Journal of the American Association for Laboratory Animal Science 48, 28–32.

Delannoy, S.M., 2005. Subdigital setae of the Tokay gecko (*Gekko gecko*): variation in form and implications for adhesion (MSc Thesis). Department of Biological Science, University of Calgary, Calgary, Alberta, CA.

Garner, A.M., Wilson, M.C., Russell, A.P., Dhinojwala, A., Niewiarowski, P.H., 2019. Going out on a limb: how investigation of the anoline adhesive system can enhance our understanding of fibrillar adhesion. Integrative and Comparative Biology 59, 61–69. https://doi.org/10.1093/icb/icz012

Ge, L., Sethi, S., Ci, L., Ajayan, P.M., Dhinojwala, A., 2007. Carbon nanotube-based synthetic gecko tapes. Proceedings of the National Academy of Sciences of the United States of America 104, 10792–10795. https://doi.org/10.1073/pnas.0703505104

Greiner, C., Arzt, E., Del Campo, A., 2009. Hierarchical gecko◻like adhesives. Advanced Materials 21, 479–482. https://doi.org/10.1002/adma.200801548

Greiner, C., del Campo, A., Arzt, E., 2007. Adhesion of bioinspired micropatterned surfaces:L effects of pillar radius, aspect ratio, and preload. Langmuir 23, 3495–3502. https://doi.org/10.1021/la0633987

Huber, G., Mantz, H., Spolenak, R., Mecke, K., Jacobs, K., Gorb, S.N., Arzt, E., 2005. Evidence for capillarity contributions to gecko adhesion from single spatula nanomechanical measurements. Proceedings of the National Academy of Sciences of the United States of America 102, 16293–16296.

Irschick, D.J., Austin, C.C., Petren, K., Fisher, R.N., Losos, J.B., Ellers, O., 1996. A comparative analysis of clinging ability among pad◻bearing lizards. Biological Journal of the Linnean Society 59, 21–35.

Johnson, K.L., Kendall, K., Roberts, A.D., 1971. Surface energy and the contact of elastic solids. Proceedings of the Royal Society of London A: Mathematical, Physical and Engineering Sciences 324, 301–313.

Johnson, M.K., Russell, A.P., 2009. Configuration of the setal fields of *Rhoptropus* (Gekkota: Gekkonidae): functional, evolutionary, ecological and phylogenetic implications of observed pattern. Journal of Anatomy 214, 937–955. https://doi.org/10.1111/j.1469-7580.2009.01075.x

Kendall, K., 1975. Thin-film peeling-the elastic term. Journal of Physics D: Applied Physics 8, 1449–1452.

Khanafer, K., Duprey, A., Schlicht, M., Berguer, R., 2008. Effects of strain rate, mixing ratio, and stress–strain definition on the mechanical behavior of the polydimethylsiloxane (PDMS) material as related to its biological applications. Biomedical Microdevices 11, 503. https://doi.org/10.1007/s10544-008-9256-6

Losos, J.B., 2011. Lizards in an Evolutionary Tree: Ecology and Adaptive Radiation of Anoles. University of California Press, Berkeley, California.

Losos, J.B., 1994. Integrative approaches to evolutionary ecology: *Anolis* lizards as model systems. Annual Review of Ecology and Systematics 25, 467–493.

Maderson, P.F.A., 1970. Lizard glands and lizard hands: models for evolutionary study. Forma et Functio 3, 179–204.

Murphy, M.P., Kim, S., Sitti, M., 2009. Enhanced adhesion by gecko-inspired hierarchical fibrillar adhesives. ACS Applied Materials & Interfaces 1, 849–855.

Niewiarowski, P.H., Lopez, S., Ge, L., Hagan, E., Dhinojwala, A., 2008. Sticky gecko feet: the role of temperature and humidity. PLoS ONE 3. https://doi.org/10.1371/journal.pone.0002192

Niewiarowski, P.H., Stark, A.Y., Dhinojwala, A., 2016. Sticking to the story: outstanding challenges in gecko-inspired adhesives. The Journal of Experimental Biology 219, 912–919. https://doi.org/10.1242/jeb.080085

Peattie, A.M., Full, R.J., 2007. Phylogenetic analysis of the scaling of wet and dry biological fibrillar adhesives. Proceedings of the National Academy of Sciences 104, 18595–18600.

Persson, B.N.J., 2003. On the mechanism of adhesion in biological systems. The Journal of Chemical Physics 118, 7614–7621.

Peterson, J.A., 1983. The evolution of the subdigital pad in *Anolis*. I. Comparisons among the anoline genera, in: Advances in Herpetology and Evolutionary Biology: Essays in Honor of Ernest E. Williams. Museum of Comparative Zoology, Harvard University, Cambridge, MA USA.

Ruibal, R., Ernst, V., 1965. The structure of the digital setae of lizards. Journal of Morphology 117, 271–293.

Russell, A.P., 2017. The structure of anoline (Reptilia: Dactyloidae: *Anolis*) toe pads in relation to substratum conformity. Acta Zoologica 98, 300–309.

Russell, A.P., Bels, V., 2001. Digital hyperextension in *Anolis sagrei*. Herpetologica 57, 58–65.

Russell, A.P., Gamble, T., 2019. Evolution of the Gekkotan Adhesive System: Does Digit Anatomy Point to One or More Origins? Integrative and Comparative Biology 59, 131–147. https://doi.org/10.1093/icb/icz006

Russell, A.P., Johnson, M.K., 2014. Between a rock and a soft place: microtopography of the locomotor substrate and the morphology of the setal fields of Namibian day geckos (Gekkota: Gekkonidae: *Rhoptropus*). Acta Zoologica 95, 299–318. https://doi.org/10.1111/azo.12028

Russell, A.P., Johnson, M.K., 2007. Real-world challenges to, and capabilities of, the gekkotan adhesive system: contrasting the rough and the smooth. Canadian Journal of Zoology 85, 1228–1238. https://doi.org/10.1139/Z07-103

Russell, A.P., Johnson, M.K., Delannoy, S.M., 2007. Insights from studies of gecko-inspired adhesion and their impact on our understanding of the evolution of the gekkotan adhesive system. Journal of Adhesion Science and Technology 21, 1119–1143. https://doi.org/10.1163/156856107782328371

Spolenak, R., Gorb, S., Arzt, E., 2005. Adhesion design maps for bio-inspired attachment systems. Acta Biomaterialia 1, 5–13. https://doi.org/10.1016/j.actbio.2004.08.004

Stork, N.E., 1983. A comparison of the adhesive setae on the feet of lizards and arthropods. Journal of Natural History 17, 829–835.

Williams, E.E., Peterson, J.A., 1982. Convergent and alternative designs in the digital adhesive pads of scincid lizards. Science 215, 1509–1511.

Yao, H., Gao, H., 2006. Mechanics of robust and releasable adhesion in biology: Bottom–up designed hierarchical structures of gecko. Journal of the Mechanics and Physics of Solids 54, 1120–1146.

