## Supporting Information for "The same but different: setal arrays of anoles and geckos indicate alternative approaches to achieving similar adhesive effectiveness"

### The Concept of Contact Splitting

The concept of contact splitting is argued to be one of the primary benefits of hierarchy in a fibrillar system, where an adhesive area is broken up into separate contact points (Arzt et al., 2003). An increased number of small contacts improves adhesion relative to one large contact. Arzt et al. (2003) applied the Johnson-Kendall-Roberts (JKR) contact theory of elastic solids to demonstrate this concept. The critical pull-off force ($F_C$) of a hemisphere in contact with a planar surface can be written as

$$F_C = \frac{3}{2}\pi R\gamma$$

*SI Equation 1*

where $R$ is equivalent to the radius of the contact and $\gamma$ is equivalent to the work of adhesion. If the contact point (e.g., an adhesive pad), however, is split into $n$ smaller contacts (e.g., setae) with radii $r$ ($r = R/\sqrt{n}$), the JKR equation can be modified to estimate the increased critical pull-off force ($F_C'$) resulting from contact splitting,

$$F_C' = \sqrt{n} \cdot F_C$$

*SI Equation 2*

Thus, it follows that the increased subdivision of an adhesive contact into smaller and more numerous contact points directly results in an increase in adhesive force.

### Effective Modulus of Tapered Setal Arrays

The bending stiffness of a non-tapered seta ($k$) can be approximated as

$$k = \frac{3E\pi R^4}{4L^3}$$

*SI Equation 3*

where $E$ is equivalent to the elastic modulus of β-keratin (~1 GPa), $R$ is equivalent to the radius of the setal stalk, and $L$ is equivalent to the setal length. Autumn et al. (2006) estimated the elastic modulus ($E_{eff}$) of gecko setal arrays (non-tapered arrays) to be:

$$E_{eff} = k \cdot L \cdot \frac{D \sin(\theta)}{(\cos\theta)^2 \cdot [1 \pm \mu \tan(\theta)]}$$

*SI Equation 4*

where $D$ is equivalent to setal density, $\theta$ is equivalent to the setal resting angle, and $\mu$ is equivalent to the setal coefficient of friction (assumed to 0.25; Autumn et al., 2006, Johnson and Russell, 2009).

As demonstrated in the manuscript, the effective bending stiffness of a tapered seta ($k_{eff}$) can be approximated by multiplying the bending stiffness of a non-tapered seta ($k$) by what we call the tapering ratio ($t$), which is simply the ratio of the apex radius ($R_a$) to the base radius ($R_b$) (Caliaro et al., 2013):

$$k_{eff} = k \cdot t = \frac{3\pi R_b^4 E}{4L^3} \cdot \frac{R_a}{R_b}$$

*SI Equation 5*

Thus, by exchanging the $k$ of a non-tapered seta with the $k_{eff}$ of a tapered seta, the effective elastic modulus of tapered setal arrays can be approximated as:

$$E_{eff} = k_{eff} \cdot L \cdot \frac{D \sin(\theta)}{(\cos\theta)^2 \cdot [1 \pm \mu \tan(\theta)]}$$

*SI Equation 6*

**Supplementary Figures**

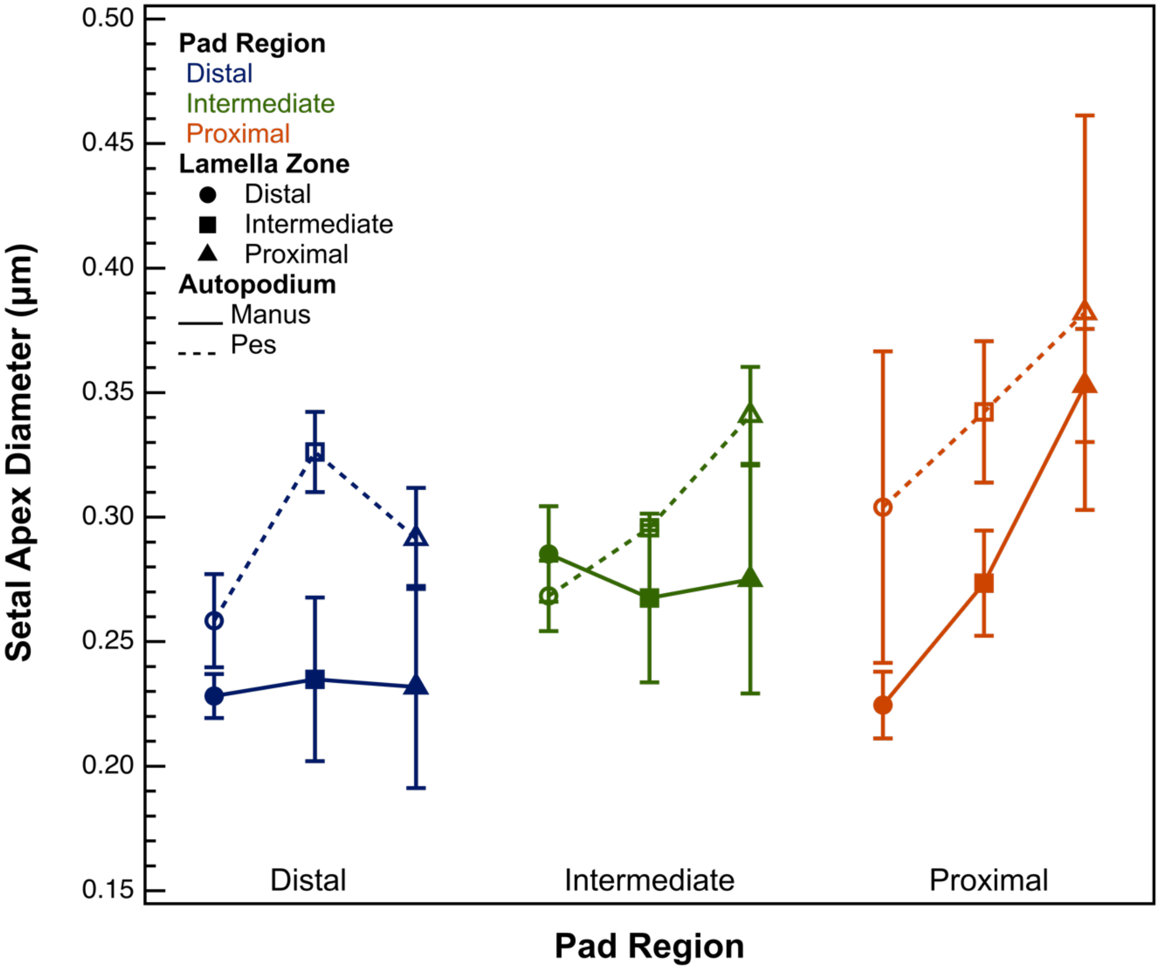


**SI Figure 1.** Mean setal apex diameter as a function of pad zone, lamella zone, and limb appendage. Mean setal apex diameter was significantly greater in the pes compared to the manus ($P < 0.0001$). In both the manus and the pes, mean setal apex diameter was significantly reduced in distal pad regions compared to proximal ones, but only in the proximal lamella zone ($P = 0.0003$). In the proximal pad region, setal apex diameter was significantly lower in distal lamella zones compared to proximal lamella zones ($P = 0.0003$). There were no other significant differences in mean setal apex diameter between different pad regions and/or lamella zones.

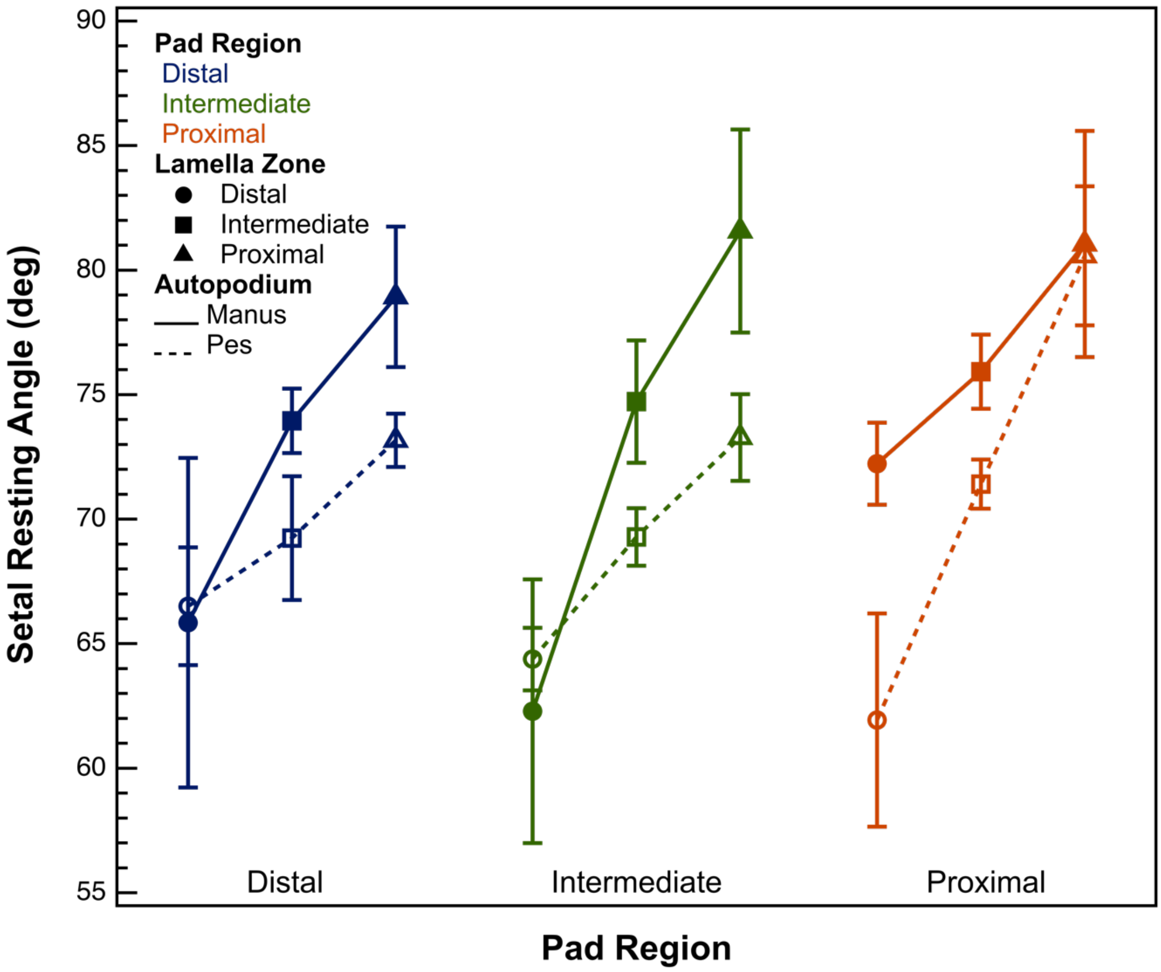


**SI Figure 2.** Mean setal resting angle as a function of pad region, lamella zone, and limb appendage. For both the manus and pes, mean setal resting angle was significantly lower in the distal pad region compared to the proximal region ($P = 0.01$), and there were no significant differences in mean setal resting angle between the setae of the intermediate pad region compared to either the distal or proximal pad region (intermediate vs. distal: $P = 0.329$; intermediate vs. proximal: $P = 0.248$). Mean setal resting angle significantly decreased proximodistally along all lamella zones on both the manus and pes (all pairwise comparisons $P < 0.05$).

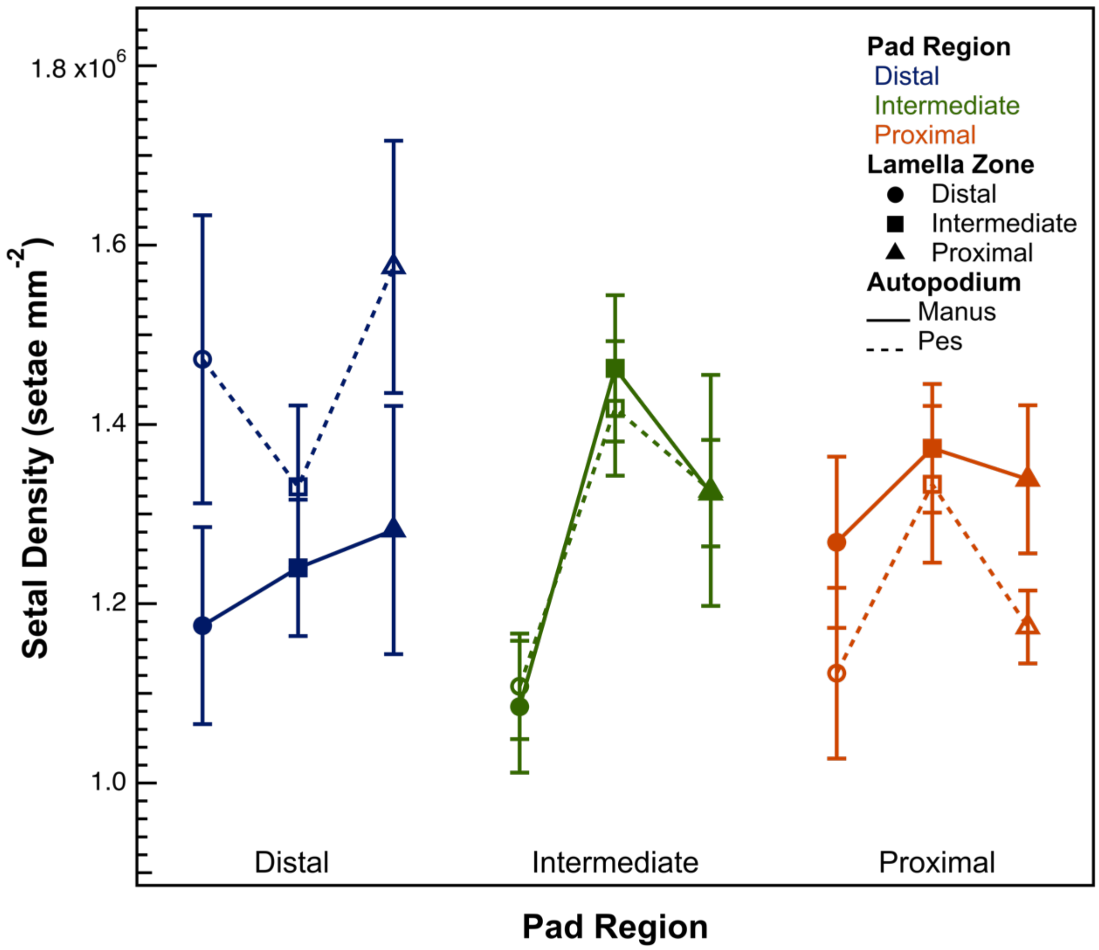


**SI Figure 3.** Mean setal density as a function of pad region, lamella zone, and limb appendage. An ANOVA revealed that mean setal density did not significantly vary across pad regions, lamella zones, or limb appendages (whole model P = 0.11).

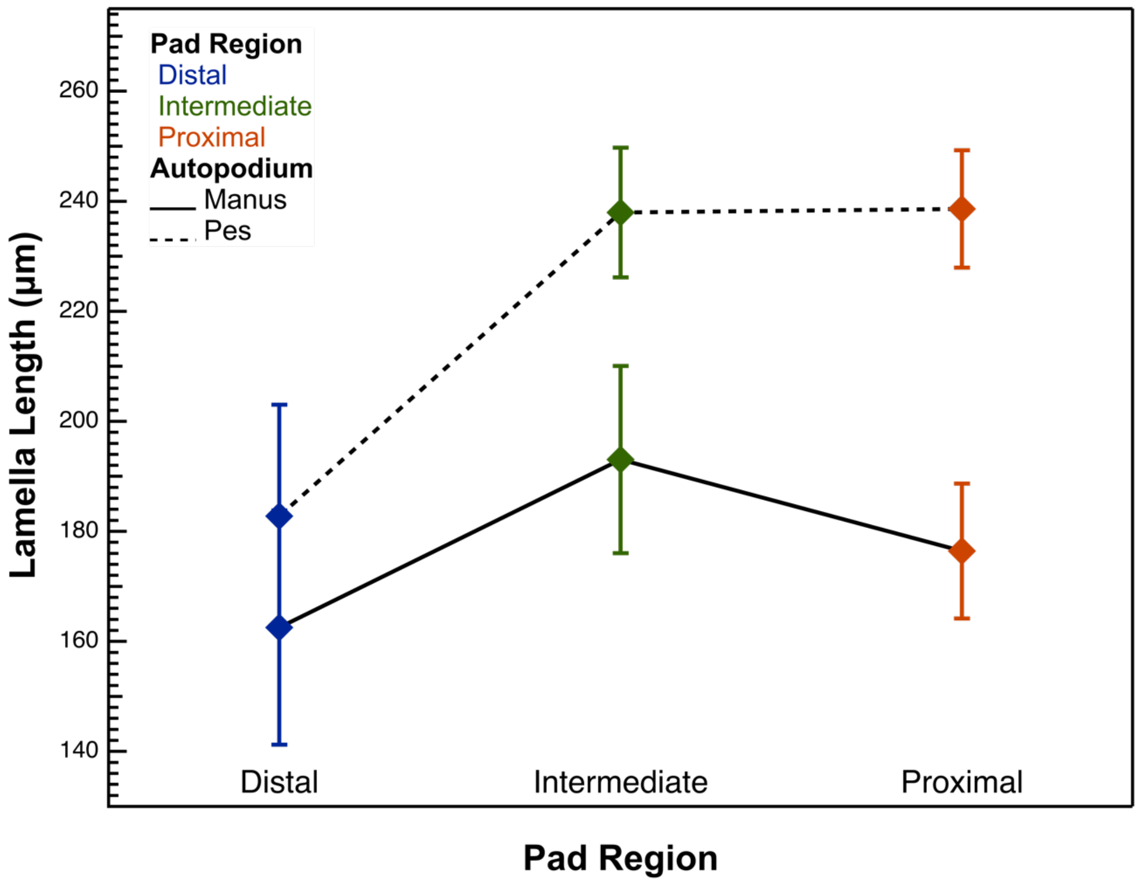


**SI Figure 4.** Mean lamella length as a function of pad region and limb appendage. The lamellae from the pes were significantly longer than those from the manus (P = 0.0009). On both the manus and the pes, lamellae from the distal regions of the pad were significantly longer than those from the intermediate and proximal regions of the pad (distal vs. intermediate: P = 0.033; distal vs. proximal: P = 0.022). Lamellae from the intermediate and proximal regions of the pad were not significantly different in length on both the manus and the pes (P = 0.985).

### Supplementary Tables

| Autopodium | Pad Region | Lamella Zone | Setal Length (µm) | Setal Base Diameter (µm) |
|---|---|---|---|---|
| Manus | Proximal | Proximal | 15.004 ± 1.418 | 0.685 ± 0.023 |
| | | Intermediate | 18.357 ± 0.604 | 0.626 ± 0.021 |
| | | Distal | 16.676 ± 0.967 | 0.531 ± 0.013 |
| | Intermediate | Proximal | 18.821 ± 0.869 | 0.667 ± 0.046 |
| | | Intermediate | 21.357 ± 0.976 | 0.619 ± 0.034 |
| | | Distal | 19.591 ± 1.85 | 0.546 ± 0.019 |
| | Distal | Proximal | 18.219 ± 1.183 | 0.626 ± 0.041 |
| | | Intermediate | 21.819 ± 0.285 | 0.598 ± 0.033 |
| | | Distal | 19.844 ± 0.918 | 0.534 ± 0.009 |
| Pes | Proximal | Proximal | 17.714 ± 1.728 | 0.738 ± 0.079 |
| | | Intermediate | 21.382 ± 0.427 | 0.595 ± 0.028 |
| | | Distal | 16.24 ± 0.518 | 0.562 ± 0.063 |
| | Intermediate | Proximal | 18.826 ± 1.423 | 0.792 ± 0.019 |
| | | Intermediate | 26.462 ± 0.739 | 0.617 ± 0.001 |
| | | Distal | 16.811 ± 0.769 | 0.53 ± 0.014 |
| | Distal | Proximal | 18.535 ± 0.836 | 0.596 ± 0.02 |
| | | Intermediate | 23.697 ± 0.837 | 0.581 ± 0.016 |
| | | Distal | 21.152 ± 0.616 | 0.499 ± 0.019 |

**SI Table 1.** Setal length and base diameter as a function of autopodium, pad region, and lamella zone. Values are mean ± 1 sem.

| Autopodium | Pad Region | Lamella Zone | Setal Apex Diameter (µm) | $k_{eff}$ (N/m) |
|---|---|---|---|---|
| Manus | Proximal | Proximal | 0.353 ± 0.036 | 0.0067 ± 0.0012 |
| | | Intermediate | 0.273 ± 0.021 | 0.0016 ± 0.0003 |
| | | Distal | 0.225 ± 0.016 | 0.0013 ± 0.0002 |
| | Intermediate | Proximal | 0.275 ± 0.018 | 0.0027 ± 0.0004 |
| | | Intermediate | 0.268 ± 0.018 | 0.0009 ± 0.0002 |
| | | Distal | 0.285 ± 0.03 | 0.0013 ± 0.0002 |
| | Distal | Proximal | 0.232 ± 0.013 | 0.002 ± 0.0002 |
| | | Intermediate | 0.235 ± 0.021 | 0.0006 ± 0.0002 |
| | | Distal | 0.228 ± 0.017 | 0.0008 ± 0.0001 |
| Pes | Proximal | Proximal | 0.382 ± 0.017 | 0.0064 ± 0.0028 |
| | | Intermediate | 0.342 ± 0.016 | 0.0012 ± 0.0002 |
| | | Distal | 0.304 ± 0.018 | 0.0016 ± 0.0003 |
| | Intermediate | Proximal | 0.341 ± 0.025 | 0.0047 ± 0.0004 |
| | | Intermediate | 0.296 ± 0.058 | 0.0006 ± 0.0001 |
| | | Distal | 0.268 ± 0.002 | 0.0015 ± 0.0003 |
| | Distal | Proximal | 0.291 ± 0.024 | 0.0017 ± 0.0005 |
| | | Intermediate | 0.326 ± 0.008 | 0.0007 ± 0.0001 |
| | | Distal | 0.258 ± 0.016 | 0.0005 ± 0.0001 |

**SI Table 2.** Setal apex diameter and effective setal bending modulus ($k_{eff}$) as a function of autopodium, pad region, and lamella zone. Values are mean ± 1 sem.

| Autopodium | Pad Region | Lamella Zone | Setal Resting Angle (deg) | Setal Density (setae/mm$^2$) |
|---|---|---|---|---|
| Manus | Proximal | Proximal | 81.046 ± 4.539 | 1338638 ± 82655 |
| | | Intermediate | 75.924 ± 1.492 | 1373368 ± 71648 |
| | | Distal | 72.228 ± 1.646 | 1268614 ± 95530 |
| | Intermediate | Proximal | 81.57 ± 4.073 | 1323341 ± 59373 |
| | | Intermediate | 74.721 ± 2.458 | 1462429 ± 81681 |
| | | Distal | 62.287 ± 5.292 | 1085207 ± 73466 |
| | Distal | Proximal | 78.929 ± 2.819 | 1282295 ± 138518 |
| | | Intermediate | 73.95 ± 1.297 | 1240006 ± 76007 |
| | | Distal | 65.841 ± 6.62 | 1175601 ± 109888 |
| Pes | Proximal | Proximal | 80.571 ± 2.795 | 1174121 ± 40644 |
| | | Intermediate | 71.41 ± 0.982 | 1333122 ± 87423 |
| | | Distal | 61.93 ± 4.278 | 1122610 ± 95103 |
| | Intermediate | Proximal | 73.276 ± 1.744 | 1326320 ± 128760 |
| | | Intermediate | 69.289 ± 1.158 | 1417921 ± 75124 |
| | | Distal | 64.385 ± 1.258 | 1107793 ± 58815 |
| | Distal | Proximal | 73.17 ± 1.066 | 1575738 ± 140739 |
| | | Intermediate | 69.236 ± 2.483 | 1330401 ± 90680 |
| | | Distal | 66.5 ± 2.362 | 1472648 ± 160704 |

**SI Table 3.** Setal resting angle and setal density as a function of autopodium, pad region, lamella zone. Values are ± 1 sem.

| Autopodium | Pad Region | Lamella Length (µm) |
|---|---|---|
| Manus | Proximal | 176.419 ± 12.252 |
| | Intermediate | 193.022 ± 17 |
| | Distal | 162.507 ± 21.28 |
| Pes | Proximal | 238.607 ± 10.651 |
| | Intermediate | 237.956 ± 11.778 |
| | Distal | 182.763 ± 20.273 |

**SI Table 4.** Lamella length as a function of autopodium and pad region. Values are mean ± 1 sem.

| Source | Setal Length | | | | Setal Apex Diameter | | | |
|---|---|---|---|---|---|---|---|---|
| | DF | SS | F | P | DF | SS | F | P |
| Autopodium | 1 | 23.04 | 6.31 | 0.015* | 1 | 0.036 | 22.47 | <0.0001* |
| Pad Region | 2 | 113.89 | 15.60 | <0.0001* | 2 | 0.026 | 8.13 | 0.0009* |
| Lamella Zone | 2 | 188.09 | 25.77 | <0.0001* | 2 | 0.023 | 7.27 | 0.0018* |
| Autopodium x Lamella Zone | 2 | 24.78 | 3.39 | 0.042* | - | - | - | - |
| Pad Region x Lamella Zone | - | - | - | - | 4 | 0.021 | 3.29 | 0.018* |

**SI Table 5.** Analysis of variance tables for setal length and setal apex diameter. * indicates significant P values.

| Source | Setal Base Diameter | | | |
|---|---|---|---|---|
| | DFNum | DFDen | F | P |
| Autopodium | 1 | 45.43 | 0.01 | 0.92 |
| Pad Region | 2 | 30.68 | 3.31 | 0.05 |
| Lamella Zone | 2 | 33.97 | 23.02 | <0.0001* |

**SI Table 6.** Welch's analysis of variance table for setal base diameter. * indicates significant P values.

| Source | $k_{eff}$ | | | |
|---|---|---|---|---|
| | DF | SS | F | P |
| Autopodium | 1 | 0.11 | 0.6 | 0.44 |
| Pad Region | 2 | 8.61 | 23.07 | <0.0001* |
| Lamella Zone | 2 | 18.9 | 50.65 | <0.0001* |
| Autopodium x Lamella Zone | - | - | - | - |
| Pad Region x Lamella Zone | - | - | - | - |

**SI Table 7.** Analysis of variance tables for effective bending modulus ($k_{eff}$). * indicates significant P values.

| Source | Setal Resting Angle | | |
|---|---|---|---|
| | ChiSquare | DF | P |
| Autopodium | 1.978 | 1 | 0.16 |
| Pad Region | 6.321 | 2 | 0.042* |
| Lamella Zone | 20.065 | 2 | <0.0001* |

**SI Table 8**. Wilcoxon rank sum test table for setal resting angle. * indicates significant P values.

| Source | Lamella Length | | | |
|---|---|---|---|---|
| | DF | SS | F | P |
| Autopodium | 1 | 11306.53 | 19.10 | 0.0009* |
| Pad Region | 2 | 7160.47 | 6.05 | 0.015* |

**SI Table 9.** Analysis of variance table for the analysis of lamella length. * indicates significant P values.

*Pad Region*

| Level | - Level | p-Value |
|---|---|---|
| Distal | Proximal | <0.0001* |
| Intermediate | Proximal | 0.0002* |
| Distal | Intermediate | 0.962 |

*Lamella Zone*

| Level | - Level | p-Value |
|---|---|---|
| Intermediate | Proximal | <0.0001* |
| Intermediate | Distal | <0.0001* |
| Distal | Proximal | 0.403 |

*Autopodium x Lamella Zone*

| Level | - Level | p-Value |
|---|---|---|
| Pes,Intermediate | Manus,Proximal | <0.0001* |
| Pes,Intermediate | Pes,Proximal | <0.0001* |
| Pes,Intermediate | Pes,Distal | <0.0001* |
| Pes,Intermediate | Manus,Distal | <0.0001* |
| Manus,Intermediate | Manus,Proximal | 0.006* |
| Pes,Intermediate | Manus,Intermediate | 0.014* |

| | | |
|---|---|---|
| Manus,Intermediate | Pes,Proximal | 0.151 |
| Manus,Intermediate | Pes,Distal | 0.246 |
| Manus,Intermediate | Manus,Distal | 0.304 |
| Manus,Distal | Manus,Proximal | 0.587 |
| Pes,Distal | Manus,Proximal | 0.734 |
| Pes,Proximal | Manus,Proximal | 0.865 |
| Manus,Distal | Pes,Proximal | 0.998 |
| Pes,Distal | Pes,Proximal | 1.000 |
| Manus,Distal | Pes,Distal | 1.000 |

**SI Table 10.** Tukey HSD table for the analysis of setal length. * indicates significant P values.

*Lamella Zone*

| Level | - Level | p-Value |
|---|---|---|
| Proximal | Distal | <0.0001* |
| Intermediate | Distal | <0.0001* |
| Proximal | Intermediate | 0.015* |

**SI Table 11.** Games-Howell multiple comparisons table for the analysis of setal base diameter. * indicates significant P values.

*Pad Region*

| Level | - Level | p-Value |
|---|---|---|
| Proximal | Distal | 0.0006* |
| Intermediate | Distal | 0.099* |
| Proximal | Intermediate | 0.267 |

*Lamella Zone*

| Level | - Level | p-Value |
|---|---|---|
| Proximal | Distal | 0.001* |
| Intermediate | Distal | 0.084 |
| Proximal | Intermediate | 0.249 |

*Pad Region x Lamella Zone*

| Level | - Level | p-Value |
|---|---|---|
| Proximal,Proximal | Distal,Distal | <0.0001* |
| Proximal,Proximal | Proximal,Distal | 0.0003* |
| Proximal,Proximal | Distal,Proximal | 0.0003* |
| Proximal,Proximal | Distal,Intermediate | 0.007* |
| Proximal,Proximal | Intermediate,Distal | 0.018* |

| | | |
|---|---|---|
| Proximal,Proximal | Intermediate,Intermediate | 0.018* |
| Proximal,Intermediate | Distal,Distal | 0.088 |
| Intermediate,Proximal | Distal,Distal | 0.160 |
| Proximal,Proximal | Intermediate,Proximal | 0.187 |
| Proximal,Proximal | Proximal,Intermediate | 0.111 |
| Proximal,Intermediate | Proximal,Distal | 0.504 |
| Proximal,Intermediate | Distal,Proximal | 0.508 |
| Intermediate,Proximal | Proximal,Distal | 0.628 |
| Intermediate,Proximal | Distal,Proximal | 0.632 |
| Intermediate,Intermediate | Distal,Distal | 0.690 |
| Intermediate,Distal | Distal,Distal | 0.697 |
| Distal,Intermediate | Distal,Distal | 0.589 |
| Proximal,Intermediate | Distal,Intermediate | 0.974 |
| Intermediate,Proximal | Distal,Intermediate | 0.986 |
| Proximal,Intermediate | Intermediate,Distal | 0.986 |
| Intermediate,Proximal | Intermediate,Distal | 0.992 |
| Proximal,Intermediate | Intermediate,Intermediate | 0.987 |
| Intermediate,Proximal | Intermediate,Intermediate | 0.992 |
| Intermediate,Intermediate | Proximal,Distal | 0.991 |
| Intermediate,Intermediate | Distal,Proximal | 0.991 |
| Intermediate,Distal | Proximal,Distal | 0.991 |
| Intermediate,Distal | Distal,Proximal | 0.991 |
| Distal,Intermediate | Proximal,Distal | 0.985 |
| Distal,Intermediate | Distal,Proximal | 0.985 |
| Distal,Proximal | Distal,Distal | 0.988 |
| Proximal,Distal | Distal,Distal | 0.988 |
| Intermediate,Intermediate | Distal,Intermediate | 1.000 |
| Intermediate,Intermediate | Intermediate,Distal | 1.000 |
| Intermediate,Distal | Distal,Intermediate | 1.000 |
| Proximal,Intermediate | Intermediate,Proximal | 1.000 |
| Distal,Proximal | Proximal,Distal | 1.000 |

**SI Table 12.** Tukey HSD table for the analysis of setal apex diameter. * indicates significant P values.

| *Pad Region* | | |
|---|---|---|
| **Level** | **- Level** | **p-Value** |
| Proximal | Distal | <0.0001* |
| Intermediate | Distal | 0.002* |
| Proximal | Intermediate | 0.037* |

*Lamella Zone*

| Level | - Level | p-Value |
|---|---|---|
| Proximal | Intermediate | <0.0001* |
| Proximal | Distal | <0.0001* |
| Distal | Intermediate | 0.632 |

**SI Table 13.** Tukey HSD table for the analysis of effective bending modulus ($k_{eff}$). * indicates significant P values.

*Pad Region*

| Level | - Level | p-Value |
|---|---|---|
| Proximal | Distal | 0.010* |
| Proximal | Intermediate | 0.248 |
| Intermediate | Distal | 0.369 |

*Lamella Zone*

| Level | - Level | p-Value |
|---|---|---|
| Proximal | Distal | 0.0002* |
| Intermediate | Distal | 0.0024* |
| Proximal | Intermediate | 0.0061* |

**SI Table 14.** Nonparametric pairwise comparisons via the Wilcoxon method for the analysis of setal resting angle. * indicates significant P values against an adjusted α after a sequential Bonferonni correction.

*Pad Region*

| Level | - Level | p-Value |
|---|---|---|
| Intermediate | Distal | 0.033* |
| Proximal | Distal | 0.022* |
| Intermediate | Proximal | 0.985 |

**SI Table 15.** Tukey HSD table for the analysis of lamella length. * indicates significant P values.

### SI References

Arzt, E., Gorb, S., Spolenak, R., 2003. From micro to nano contacts in biological attachment devices. Proceedings of the National Academy of Sciences of the United States of America 100, 10603–10606. https://doi.org/10.1073/pnas.1534701100

Autumn, K., Majidi, C., Groff, R.E., Dittmore, A., Fearing, R., 2006. Effective elastic modulus of isolated gecko setal arrays. Journal of Experimental Biology 209, 3558–3568.

Caliaro, M., Schmich, F., Speck, T., Speck, O., 2013. Effect of drought stress on bending stiffness in petioles of *Caladium bicolor* (Araceae). American Journal of Botany 100, 2141–2148. https://doi.org/10.3732/ajb.1300158

Johnson, M.K., Russell, A.P., 2009. Configuration of the setal fields of *Rhoptropus* (Gekkota: Gekkonidae): functional, evolutionary, ecological and phylogenetic implications of observed pattern. Journal of Anatomy 214, 937–955.
